# Integrative multi-cohort analysis reveals consistent sex differences in gut microbiota of multiple sclerosis patients

**DOI:** 10.64898/2026.04.17.719247

**Authors:** Irene Soler-Sáez, Cristina Galiana-Roselló, Rubén Grillo-Risco, Gwen Falony, Vanja Tepavčević, Sara Vieira-Silva, Francisco García-García

## Abstract

Biological sex is a key determinant in the onset and progression of multiple diseases. In multiple sclerosis (MS), females exhibit higher disease prevalence, earlier onset, and more pronounced inflammatory activity, whereas males tend to experience a more severe neurodegenerative course, characterized by accelerated central nervous system damage and increased brain atrophy. The gut microbiome has emerged as a critical factor in MS, as its composition can either ameliorate or exacerbate disease progression. In this study, we aimed to identify reproducible sex-associated differences in gut microbial composition across independent cohorts of MS patients. Through a systematic search we identified six independent studies based on 16S rRNA gene sequencing, comprising a total of 337 samples. Despite substantial inter-study variability, sex-associated differences were more pronounced in MS patients than in healthy controls. We identified 11 microbial taxa showing significant sex-associated differences in MS, nine enriched in females and two in males. Notably, the female-enriched taxa *Eggerthella* and *Eisenbergiella* were associated with specific MS subtypes and higher disability. To facilitate the use of our findings by the scientific community, we developed a freely accessible web-based tool that provides full access to our results. Thus, in this work we identified consistent and reproducible sex differences in the gut microbiota of MS patients, highlighting the importance of incorporating sex as a critical variable in microbiome research, with potential implications for understanding disease heterogeneity in MS.

**IMPORTANCE:** Multiple sclerosis (MS) affects females and males differently, but the biological reasons behind these differences are not fully understood. One potential factor is the gut microbiome (i.e., the community of microorganisms living in our intestines) which can influence immune function and disease progression. In this study, we analyzed data from multiple independent cohorts and found consistent differences in gut microbial composition between female and male MS patients. Notably, certain bacteria were more abundant in females and were linked to more severe disease features. We also developed a freely accessible web tool where researchers can explore the complete findings in detail. Our results highlight the importance of considering sex as a key factor in microbiome research and may help guide more personalized approaches to understanding and treating MS.

## INTRODUCTION

Multiple sclerosis (MS) is a chronic immune-mediated neurodegenerative condition of the central nervous system (CNS). In this disease, peripheral immune cells trigger autoreactive responses against the myelin sheath, leading to the formation of focal plaques that cause structural and functional damage (1, 2). MS is a highly heterogeneous disease structured in distinct clinical courses - relapsing-remitting (RR)MS, primary progressive (PP)MS and secondary progressive (SP)MS - that represent a dynamic continuum between inflammatory and neurodegenerative mechanisms (3).

The gut microbiota has recently emerged as an important contributor to MS pathogenesis. Studies in animal models have determined that the disease does not develop in model organisms lacking a gut microbial community (4, 5). Moreover, fecal microbiota transplantation from MS patients triggered disease progression in the experimental autoimmune encephalomyelitis (EAE) mouse model (6, 7). These effects are thought to be partly mediated by microbial-dependent T cell regulation at the gut mucosal interface, extending to systemic level immune alterations (8). The increased permeability of the gastrointestinal and blood-brain barriers associated with MS may facilitate the translocation of bacterial components that can influence the synthesis and maintenance of the myelin sheath (9, 10). In human cohorts, multiple investigations have reported alterations in the abundance of specific taxa when comparing gut microbiomes of patients with MS to healthy controls (6, 11–21). However, these findings have not been consistent across studies (22, 23), prompting efforts to integrate results across cohorts (24, 25).

MS heterogeneity is modulated by host factors such as sex, which strongly influences MS epidemiology and clinical outcomes (26). Female patients exhibit higher disease prevalence, younger age at onset, and more pronounced inflammatory activity. Meanwhile, male patients tend to suffer from more severe neurodegenerative progression, characterized by accelerated CNS damage and increased brain atrophy (27–29). In light of these differences, distinct research approaches have explored and identified potential molecular and cellular mechanisms through which sex acts as a key biological determinant in MS pathophysiology (30–34). The contribution of sex to MS may extend beyond host-intrinsic mechanisms to include sex-specific differences in gut microbiota composition. Gut microbial communities display sex-associated differences across the human lifespan (35), being shaped by hormonal, immunological, and lifestyle-related factors that modulate host-microbiome interactions (36, 37). Sex-associated microbiota differences have been described in murine models of autoimmune diseases (e.g., type 1 diabetes (38) and primary biliary cholangitis (39)) and neurodegenerative disorders (e.g., Alzheimer’s disease (40) and Parkinson’s disease (41)). For Parkinson’s disease, this evidence has also been reported in human studies (42). Despite its description in other conditions, biological sex represents a major source of heterogeneity in MS that has not yet been systematically integrated into microbiome research.

Accordingly, this study aimed to identify sex-dependent differences in the gut microbiota composition of MS patients by integrating independent 16S rRNA sequencing datasets. By adopting a meta-analysis approach, we aimed to identify consistent sex-differential abundance patterns robust to inter-study heterogeneity. We further validated the identified patterns in the largest MS cohort to date (15), and examined whether sex-differential abundant genera are also associated with clinical features of MS as disease severity and its clinical course.

## RESULTS

### Systematic review and datasets overview

Data search results are summarized in **Figure 1A**. Ten studies met all predefined criteria for our analysis. Of these, we excluded three studies due to unexpected taxonomic composition (**Supplementary Figure 1**), and reserved one study for validation of the integrative approach. After sample-level quality control, we excluded 12 samples based on the criteria detailed in the *Material and Methods* section. Thus, we included six studies in the meta-analysis, comprising a total of 337 samples. **Table 1** depicts the characteristics of the final sample set for each study.

**Figure 1.**
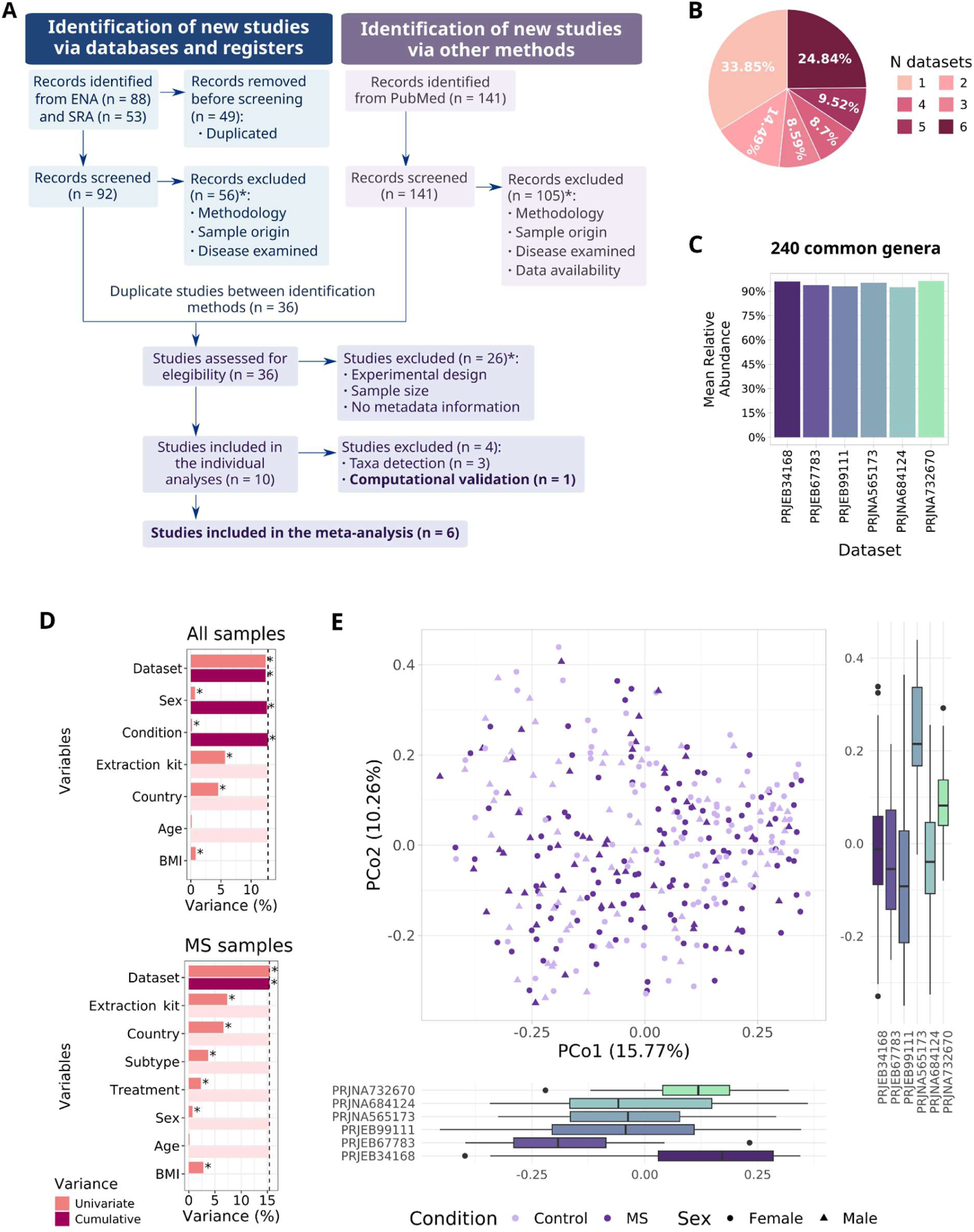
Overview of study selection and cross-dataset characterization. (A) PRISMA flow diagram summarizing the results from the study selection process. *Studies have been excluded for one or more reasons simultaneously. (B) Percentage of taxa identified according to the number of studies in which they were detected. (C) Relative abundance of taxa shared across all included datasets. (D) Microbiome compositional variation explained by reported variables in all (top) and MS (bottom) samples, either individually (univariate) or in a multivariate model (cumulative). BMI was excluded from the multivariate model due to missing data in two studies (N missing data = 143). * FDR < 0.05; dashed line: total cumulative variance. (E) Principal coordinates sample distribution, colored by condition and sex, with study-specific distributions represented as boxplots. *BMI: body mass index; ENA: European Nucleotide Archive; FDR: false discovery rate; MS: multiple sclerosis; PCo: Principal Coordinate; PRISMA: Preferred Reporting Items for Systematic Reviews and Meta-Analyses; SRA: Sequence Read Archive*.

**Table 1.** Sample characteristics by dataset included in the meta-analysis. BMI: body mass index; MS: multiple sclerosis; PMID: PubMed IDentifier; PPMS: primary progressive multiple sclerosis; RRMS: relapsing-remitting multiple sclerosis; SD: standard deviation; SPMS: secondary progressive multiple sclerosis.

|  | PRJNA684124 | PRJNA732670 | PRJEB99111 | PRJEB34168 | PRJNA565173 | PRJEB67783 |
| --- | --- | --- | --- | --- | --- | --- |
| **Sequencing platform** | Illumina MiSeq | Illumina MiSeq | Illumina MiSeq | Illumina MiSeq | Illumina MiSeq | Illumina MiSeq |
| **Sequencing read type** | Paired-end | Paired-end | Single-end | Paired-end | Paired-end | Paired-end |
| **16S hypervariable region(s)** | V3-V4 | V3-V4 | V4 | V4 | V3-V4 | V3-V4 |
| **Collection date: from - to (year)** | 2020 - 2020 | 2017 - 2019 | 2013 - 2016 | 2013 - 2017 | 2015 - 2015 | 2021 - 2022 |
| **DNA extraction kit manufacturer** | Qiagen | Qiagen | MoBio | MoBio | Qiagen | Biomes |
| **Country** | Italy | USA | USA | USA | Russia | Italy |
| **N subjects** | 30 | 56 | 113 | 65 | 30 | 43 |
| **N controls (female:male)** | 15 (7:8) | 34 (28:6) | 53 (25:28) | 39 (27:12) | 15 (9:6) | 12 (7:5) |
| **N MS (female:male)** | 15 (11:4) | 22 (17:5) | 60 (39:21) | 26 (22:4) | 15 (6:9) | 31 (19:12) |
| **Age (mean ± SD)** | 60.67 ± 13.39 | 42.91 ± 11.68 | 43.19 ± 13.06 | 43.49 ± 12.18 | 40.3 ± 15.99 | 43.73 ± 14.20 |
| **BMI (mean ± SD)** | Not reported | 26.53 ± 6.08 | Not reported | 28.19 ± 5.86 | 23.46 ± 3.25 | 23.37 ± 3.79 |
| **MS subtype** | RRMS | RRMS | RRMS | RRMS | PPMS | RRMS, SPMS |
| **Treatment** | Treated (n = 6)<br>Untreated (n = 9) | Treated (n = 16)<br>Untreated (n = 6) | Untreated (n = 60) | Untreated (n = 26) | Untreated (n = 15) | Treated (n = 30)<br>Untreated (n = 1) |
| **PMID** | 34206853 | 35472144 | 28893978 | 32743517 | 31888483 | 39402079 |

All studies recorded disease status (i.e., MS or control) and sex (i.e., female or male) information. Total samples were well balanced across conditions (168 controls and 169 MS samples), while the sex distribution skewed toward females (217 females and 120 males), consistent with the known sex-differential prevalence in MS.

The studies originated from diverse geographic regions. Age information was available for all of them, while the body mass index (BMI) was reported in four out of the six. The studies covered the three MS subtypes, and included samples from treated and untreated individuals at the time of stool collection. All data were generated using Illumina MiSeq sequencing, targeting either the V4 or the V3-V4 regions of the 16S rRNA gene.

Across all samples, we identified 965 distinct genera, being many of these study- and sample-specific. Only 24.87% (240 genera) were consistently detected across all datasets (**Figure 1B**). Importantly, these shared taxa accounted for more than 90% of the total relative abundance in every dataset, with a mean relative abundance of 94.49% (**Figure 1C**). Alpha diversity significantly differed across studies (Kruskal-Wallis test, *p* = 3.55 × 10⁻⁵), with lower diversity observed in datasets targeting the V4 region than those targeting V3-V4 (**Supplementary Figure 2A**). However, any significant differences were observed when stratifying samples by condition and sex (Kruskal-Wallis test, *p* = 0.21; **Supplementary Figure 2B**).

We next quantified the contribution of reported variables to microbiota compositional variation (**Figure 1D, top**). All variables except *age* (adjusted *p* = 0.058) were significant in univariate models, with the *dataset* variable being the main contributor to variation in microbial composition. In the multivariate model, *dataset*, *condition*, and *sex* remained significant, jointly explaining 12.8% of non-redundant variance. Consistently, principal coordinate analysis (PCoA) captured the main patterns of variation in microbiota composition (**Figure 1E**). We next restricted the analysis to MS samples that incorporated disease-specific clinical features (**Figure 1D, bottom**). The *dataset* variable remained the dominant source of variation (R² = 12.4% across all samples and 15.5% in MS samples), with no multivariate model improving upon its explanatory power.

After identifying the *dataset* variable as the main contributor to variation in the combined analysis, we explored potential sources of variability within each dataset (**Supplementary Table 1**). Most variables did not display significance; however, *condition* represented the most recurrent factor, reaching significance in three out of the six datasets explaining approximately 1-3% of the variance.

### Identification of sex-differential abundant genera in individual studies

Although *dataset* captured the major source of variation, *condition* and *sex* still emerged as significant contributors to host-related microbiome taxonomic variation. We next sought to identify sex-associated abundance patterns by condition within each dataset, preserving between-dataset variability for the subsequent meta-analysis.

Sex differential patterns were assessed for controls (control females *vs.* control males) and MS (MS females *vs.* MS males) populations separately. **Supplementary Figure 3** provides complete results for both comparisons. We observed substantial heterogeneity in genus-level abundance patterns across datasets in both comparisons, with no taxa reaching significance after FDR correction (adjusted *p* < 0.05). In the controls comparison, fewer taxa exhibited significant trends (*p* < 0.05, n = 17) than in the MS comparison (*p* < 0.05, n = 34), suggesting pathophysiology increases the gut microbiome sex differences. Meanwhile, sex differential patterns in MS comparison were qualitatively more consistent. Specifically, 40 taxa exhibited a tendency toward higher abundance in females (taxa highlighted in red) or in males (taxa highlighted in blue) in the majority of the datasets (**Figure 2**).

**Figure 2.**
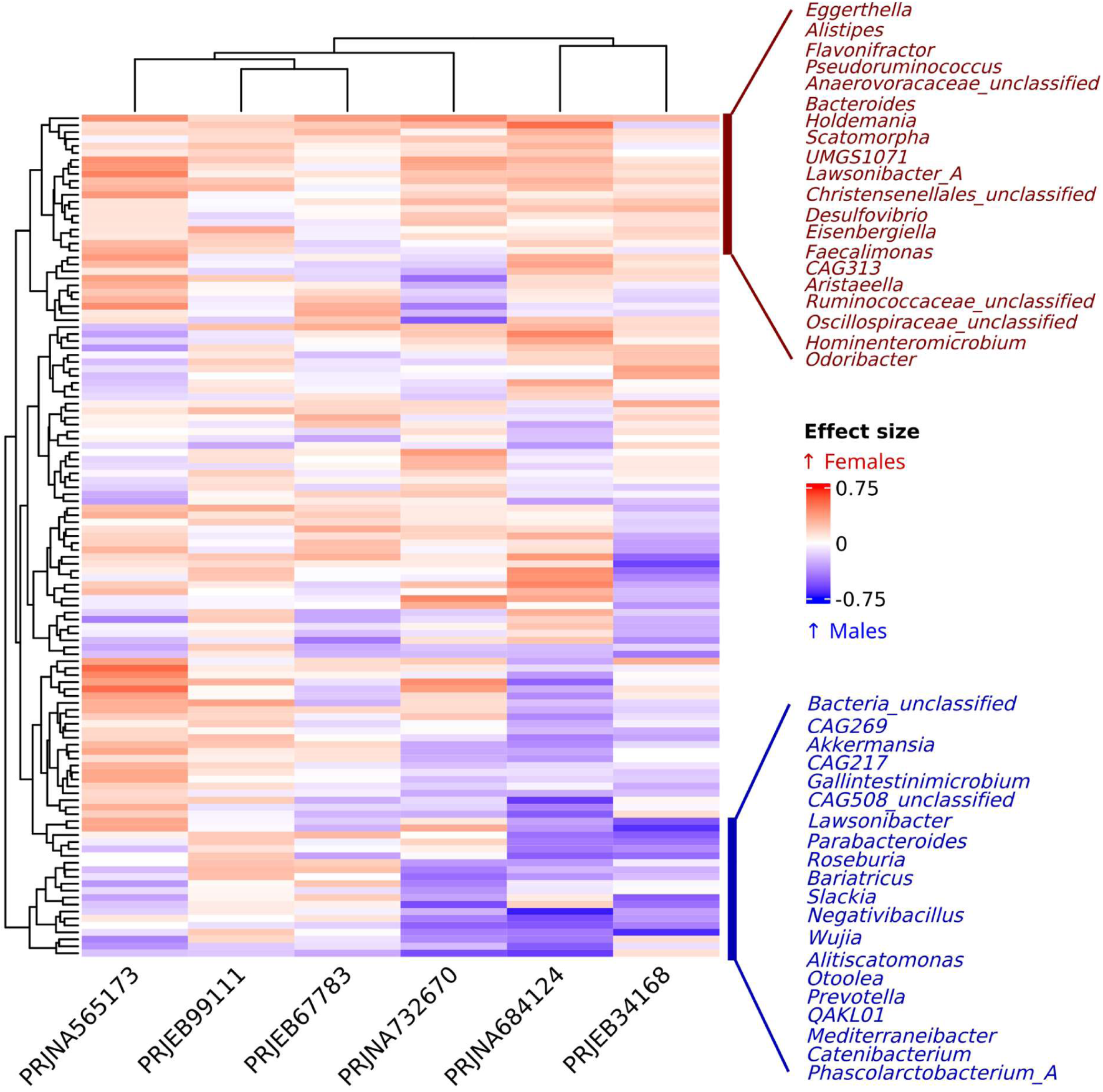
Sex-differential abundance patterns by individual datasets from MS population. Columns (datasets) and rows (genera) clustered based on hierarchical clustering. The color indicates the direction of the effect size (red: higher abundance in females, blue: higher abundance in males). Clusters with the most sex-differential consistent patterns are highlighted: female-enriched (red) and male-enriched (blue).

Taken together, although the microbiome compositional data were heterogeneous across individual studies, different taxa-most notably in the MS comparison-exhibited consistent sex differential patterns, potentially reflecting changes robust to study-specific technical and biological factors.

### Meta-analysis-based integration uncovered consistent sex-differential patterns in multiple sclerosis

To statistically assess the robustness of the sex-associated patterns observed in the individual datasets, we performed a random-effects meta-analysis for the controls and MS comparisons. No genera showed significant sex-differential abundance in healthy individuals, although eight exhibited significant trends prior to multiple testing correction (*p* < 0.05), with four more abundant in females (*Coprobacter*, *Agathobaculum*, *Eisenbergiella*, and *Evtepia*) and four more abundant in males (*Gemmiger*, *Limivivens*, *Slackia*, and *CAG217*).

Strikingly, we observed 11 genera significantly differentially abundant in the MS comparison after FDR correction (adjusted *p* < 0.05), with nine genera more abundant in females (*Desulfovibrio*, *Eggerthella*, *Eisenbergiella*, *Flavonifractor*, *Holdemania*, *Lawsonibacter*, *Pseudoruminococcus*, *Scatomorpha*, and *UMGS1071*) and two more abundant in males (*Catenibacterium* and *Prevotella*). Influence analyses revealed that no more than two of the six studies substantially contribute the combined effect size (i.e., the majority of studies contribute in a balanced manner to the consensus effect size and did not show evidence of publication bias or small-study effects). Funnel plots displayed the expected symmetric, inverted funnel shape. Together, these results support the reliability of the meta-analysis outcomes. As an illustrative example, we report the result for *Eggerthella* in **Figure 3A**, along with the corresponding influence and funnel plots in **Supplementary Figure 4A. Supplementary Figures 5-7** provide the results for the remaining 10 significant genera.

**Figure 3.**
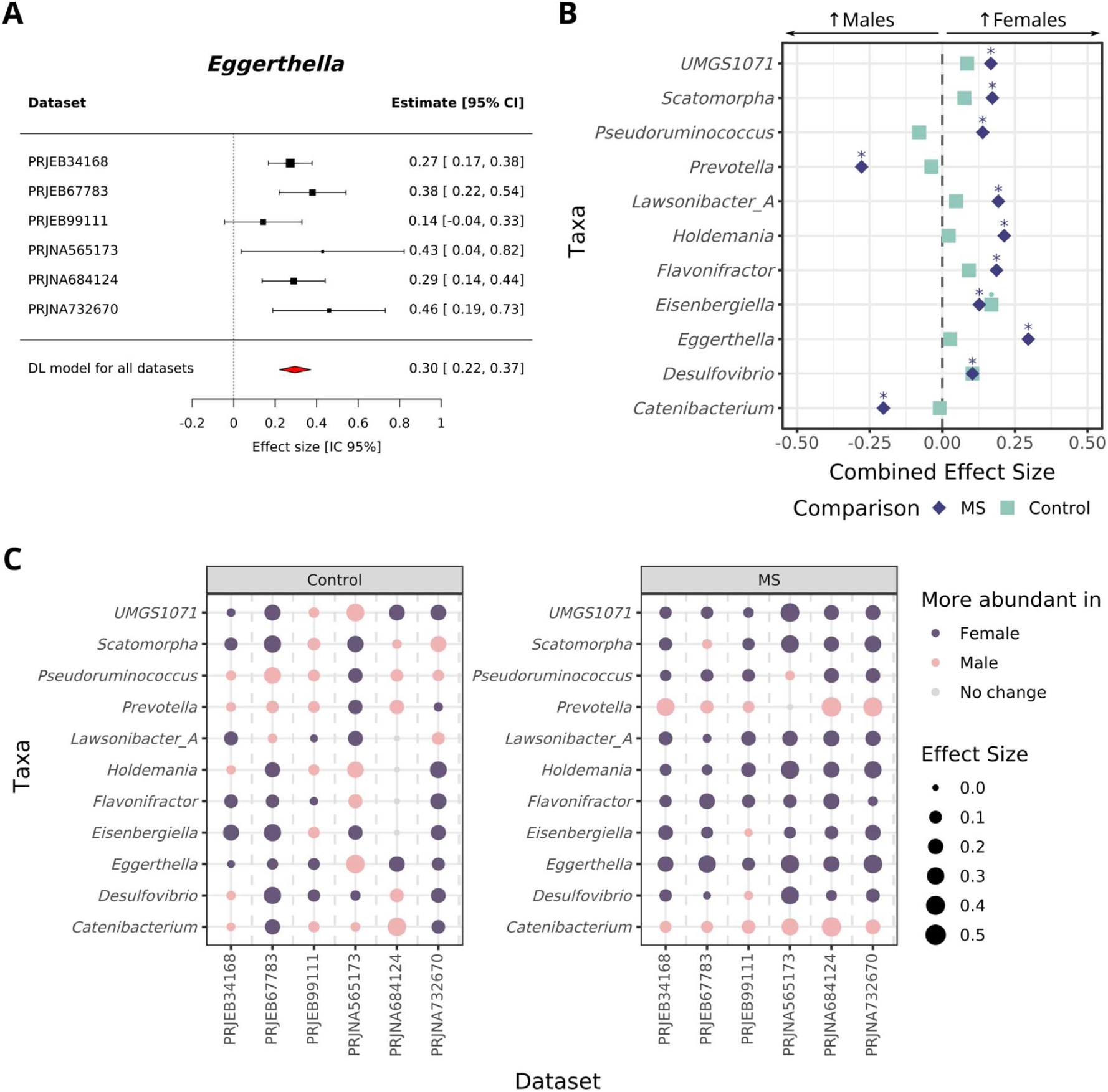
Integrative assessment of sex-associated microbial differences. (A) Forest plot for *Eggerthella* in the MS females *vs.* MS males comparison. Squares represent individual effect sizes, and horizontal lines indicate the 95% confidence intervals. The red diamond denotes the combined effect size and its confidence interval. Positive values: higher abundance in MS females; negative values: higher abundance in MS males. (B) Sex-differential significant taxa in MS. X-axis indicates the combined effect size of the comparison of females *vs.* males in controls (squares) and MS (diamonds) groups. * Adjusted p < 0.05; ⚫ p < 0.05. (C) Individual effect sizes or taxa identified as significant in the meta-analysis for controls (left) and MS (right) comparisons. *CI: Confidence Interval; DL: DerSimonianLaird; MS: multiple sclerosis*.

The significant differences identified in the MS comparison were mainly sex-associated patterns already present in healthy individuals, rather than by opposite sex-specific trends between health and disease states (**Figure 3B**). Thus, the majority of taxa showed effect sizes with the same direction in the controls as in the MS comparison, although with smaller magnitudes in the former, with both greater heterogeneity across individual studies and overall weaker effects (**Figure 3C, left**). By contrast, the MS comparison showed more consistent patterns with larger effect sizes across datasets, with six of the eleven taxa displaying concordant effects in all studies, and the remaining in five out of the six (**Figure 3C, right**). Both the heterogeneity in the controls comparison and the consistency of results in the MS comparisons suggest that MS pathophysiology increases sex-differences in gut microbiome composition.

### Validation of the identified genera in an independent cohort

To validate the findings, we analyzed the data from the iMSMS cohort (15) retrieved from the PRJEB32762 dataset. Technical and biological characteristics of this dataset are summarized in **Table 2**. This study was selected as the validation cohort due to its large sample size (N = 1,152) and the inclusion of participants from multiple geographical regions.

**Table 2.** Summary of demographic, clinical, and technical characteristics of the validation dataset. The number of samples per category is shown for categorical variables. The mean value was determined for numerical variables. *BMI: body mass index; CNS: central nervous system; EDSS: Expanded Disability Status Scale; MS: multiple sclerosis; PMID: PubMed IDentifier; PPMS: primary progressive MS; RRMS: relapsing-remitting MS; SD: standard deviation; SPMS: secondary progressive MS*.

| Variable | Total (n = 1,152) |
| --- | --- |
| Enrollment group | 1 (n = 256, recruited before October 2016), 2 (n = 896, recruited before January 2019) |
| Sequencing platform | Illumina MiSeq |
| Sequencing read type | Paired-end |
| 16S hypervariable region(s) | V4 |
| Collection date: from - to (year) | 2015 - 2019 |
| DNA extraction kit manufacturer | Qiagen |
| Recruitment site | San Francisco (n = 328), Boston (n = 84), New York (n = 118), Pittsburgh (n = 24), Buenos Aires (n = 258), Edinburgh (n = 262), San Sebastián (n = 78) |
| N controls (female:male) | 201:375 |
| N MS (female:male) | 400:176 |
| Age (mean ± SD) | 49.73 ± 12.53 |
| BMI (mean ± SD) | 26.11 ± 5.28 |
| MS subtype | RRMS (n = 437), PPMS (n = 71), SPMS (n = 68) |
| Treatment | Treated (n = 367), untreated (n = 209) |
| MS duration years (mean ± SD) | 14.21 ± 11.13 (not available = 1) |
| EDDS (mean ± SD) | 2.6 ± 2.24 |
| PMID | 36113426 |

The technical variables *enrollment group* and *recruitment site* constituted the largest known contributors to the microbial composition (**Figure 4A-B**). The *recruitment site* was also associated with significant differences in alpha diversity (**Supplementary Figure 8A-B**), in agreement with the results from the original publication (15). Therefore, we accounted for both variables in the differential abundance analyses. Beyond technical factors, MS males presented higher alpha diversity (**Supplementary Figure 8C**). The host-related biological variables recorded in this cohort-including *condition*, *BMI*, *age*, and *sex*-were incorporated into the multivariate model, together explaining 9.19% of the total variance. For MS samples, the *cohort* and *recruitment site* remained the dominant contributors to variability. In this subset, these two variables, together with the *Expanded Disability Status Scale* (*EDSS)*, *sex*, and *BMI*, explained a total of 9.21% of non-redundant variance (**Figure 4A**).

**Figure 4.**
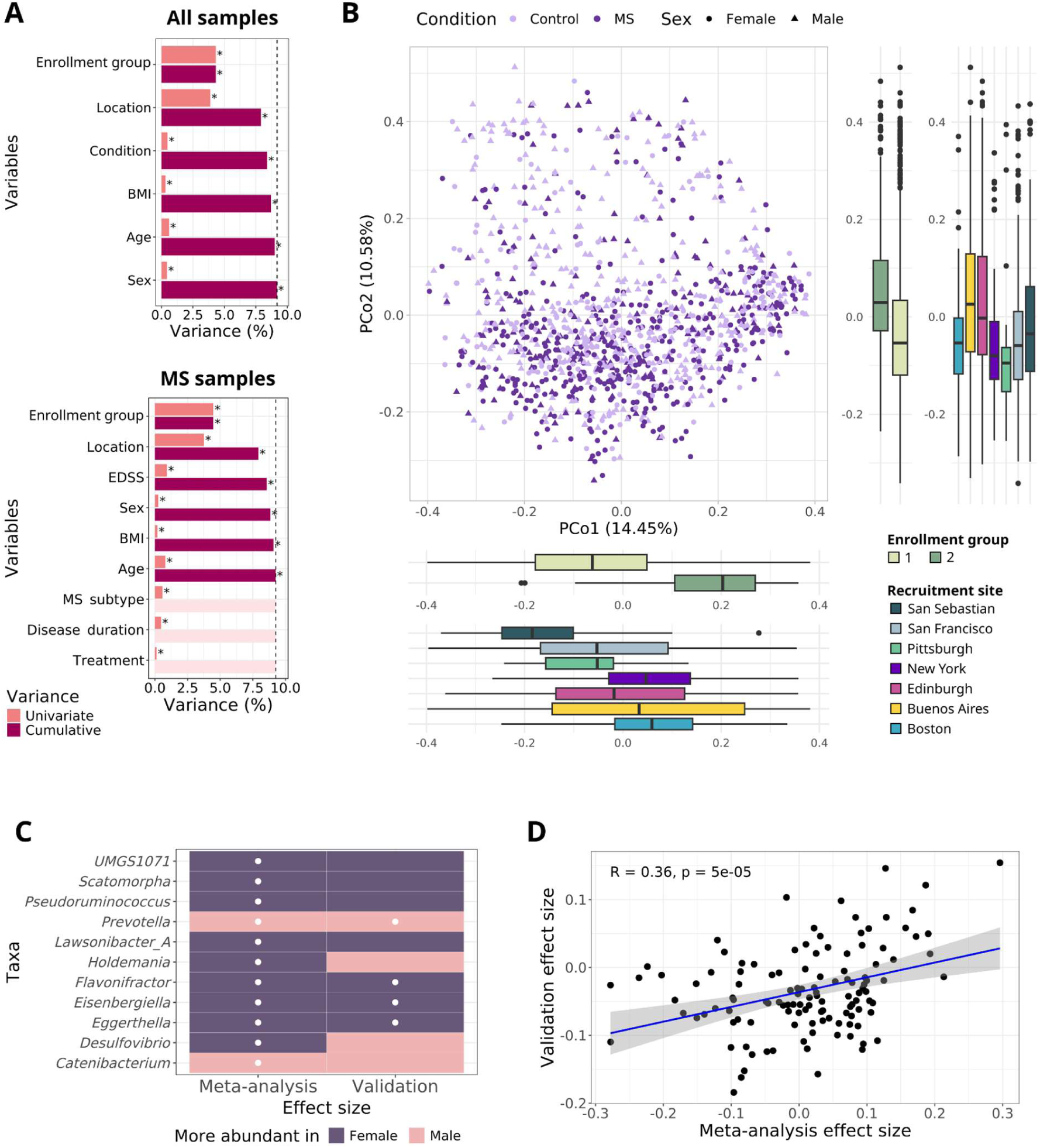
Characterization of the validation cohort and concordance of sex-differential abundance patterns in the MS comparison (MS females *vs.* MS males). (A) Microbiome compositional variation explained by reported variables in all (top) and MS (bottom) samples, either individually (univariate) or in a multivariate model (cumulative). * Adjusted p < 0.05; dashed line: total cumulative variance. (B) Principal coordinates sample distribution, colored by condition and sex. Enrollment group- and recruitment site-specific distributions are represented as boxplots. (C) Direction of effect (higher abundance in MS females vs. MS males) for taxa identified as significant in the meta-analysis, compared with corresponding results in the validation cohort. White dot: adjusted p < 0.05. (D) Pearson correlation between the combined effect sizes from the meta-analysis and those obtained in the validation cohort. Blue line: linear regression fit; shaded areas: standard error. *BMI: body mass index, EDSS: Expanded Disability Status Scale, MS: multiple sclerosis*.

The significant sex differences identified in the meta-analysis approach were mostly replicated, with 82% of the taxa (9 out of 11) showing concordant directionality between the meta-analysis and the validation dataset (**Figure 4C**). Among these nine, four taxa reached statistical significance in the validation cohort even after FDR correction. Namely, *Eggerthella*, *Eisenbergiella*, and *Flavonifractor* were more abundant in MS females, whereas *Prevotella* was more abundant in MS males. Correlating the effect sizes of the sex associations identified from the meta-analysis consensus with the results obtained in the validation cohort resulted in a positive correlation (ρ = 0.36, p < 0.05) (**Figure 4D**), supporting the reproducibility of sex-associated gut microbiome patterns in MS despite substantial inter-study variability.

### The sex-differentially abundant genera *Eggerthella* and *Eisenbergiella* are associated with MS clinical features

We next assessed whether the four validated sex-differential abundant genera with strongest statistical support were associated with MS clinical features using the phenotypic information from the iMSMS cohort from PRJEB32762 dataset (validation dataset), including MS subtype, EDSS, disease duration, and current treatment. Significant associations were identified for the female-associated *Eggerthella* and *Eisenbergiella*, but not for *Flavonifractor* nor the male-associated *Prevotella*.

*Eggerthella* was identified as more abundant in the SPMS subtype, corresponding to a more advanced disease stage after RRMS (**Figure 5A**). Additionally, its abundance was slightly positively associated with disease duration (ρ = 0.10, *p* = 0.01) and age (ρ = 0.10, *p* = 0.01). As disease duration and age were themselves moderately correlated in MS samples (ρ = 0.57, *p* < 2.2e-16), we evaluated whether this age association reflected disease-related effects. Notably, we observed no association between *Eggerthella* abundance and age was observed in the control group (ρ = 0.04, p = 0.33), suggesting that the observed associations could be linked to MS-related processes rather than to age alone.

**Figure 5.**
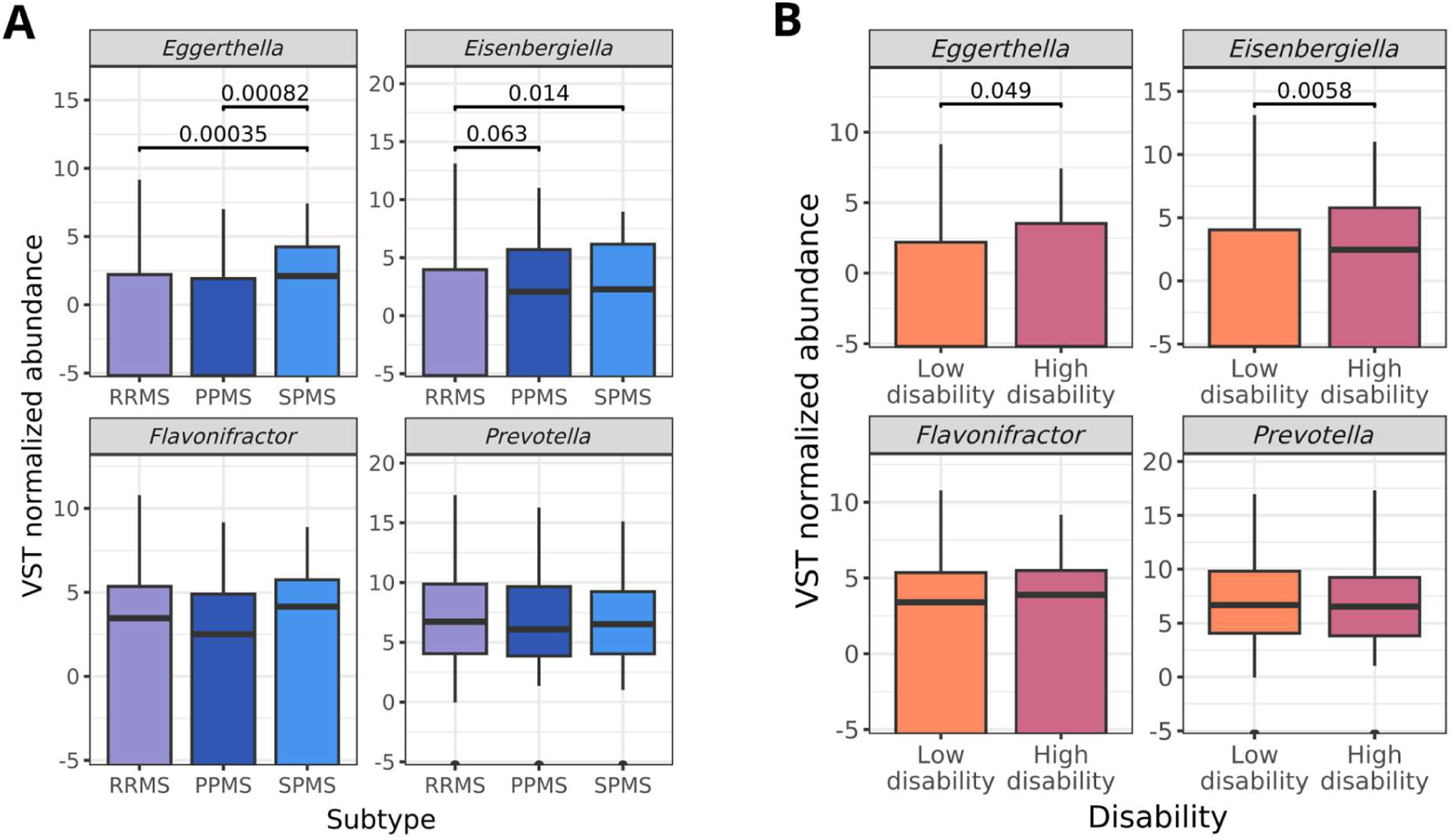
Normalized abundance of sex-associated taxa in multiple sclerosis stratified by (A) disease subtype and (B) degree of disability. Values indicate adjusted p-values (A) from Dunn’s post hoc test (after *p* < 0.05 in Kruskal–Wallis test) and (B) Wilcoxon rank-sum test. *PPMS: primary progressive multiple sclerosis; RRMS: relapsing-remitting multiple sclerosis; SPMS: secondary progressive multiple sclerosis; VST: variance stabilizing transformation*.

Conversely, *Eisenbergiella* was found to be more abundant in progressive forms of multiple sclerosis (PPMS and SPMS) than in RRMS (**Figure 5A**). Its abundance was slightly positively correlated with disease duration (ρ = 0.09, *p* = 0.03) and age (ρ = 0.13, *p* = 1.4 × 10⁻³). Unlike *Eggerthella*, the age association was replicated in the controls cohort (ρ = 0.14, *p* = 8.3 × 10⁻⁴), suggesting that age-related changes in *Eisenbergiella* are not MS-specific. Higher *Eisenbergiella* abundance was observed in patients with greater disability (EDSS > 4) relative to those with mild and low-grade disease forms (**Figure 5B**). However, patients with higher disability were significantly older (Wilcoxon test, *p* < 2.2e-16), suggesting its role in MS could be confounded by age.

*Eggerthella* and *Eisenbergiella* exhibited associations with different MS features. They were not consistently detected together across MS samples. *Eggerthella* was detected in 62.5% of samples, while *Eisenbergiella* was identified in 54%, both taxa being simultaneously detected in 23.5% of samples. In cases where both taxa were present, their abundances were positively correlated (ρ = 0.32, *p* < 0.0018) suggesting similar fitness responses to host-related factors.

None of the four validated genera showed significant differences in abundance between treated and untreated groups (**Supplementary Figure 9**), suggesting that treatment effects do not confound the observed associations. Furthermore, no clinical or demographic variable examined exhibited sex-specific differences (i.e., they present similar distributions in females and males) (**Supplementary Figure 10**).

### Web-based resource for interactive exploration of results

The complete findings from this study are available through a publicly accessible web resource (https://irsoler.shinyapps.io/metaanalisis_16S_MS/) (**Figure 6**), which provides an interactive interface organized into four tabs. The *Home* tab presents a graphical overview of the study design and objectives. The *Dataset composition* tab displays the taxonomic composition for each sample, including total counts, the number of unique taxa, rarefaction curves, and the total and relative abundance of the 15 most abundant taxa per dataset. The *Differential abundance analysis* tab allows users to explore sex-associated differential abundance results for each dataset in control and MS populations, enabling examination of metrics for specific genera and application of user-defined filters. Finally, the *Meta-analysis* tab provides interactive access to forest, funnel, and influence plots for selected genera and comparisons, along with summary tables containing the corresponding results.

**Figure 6.**
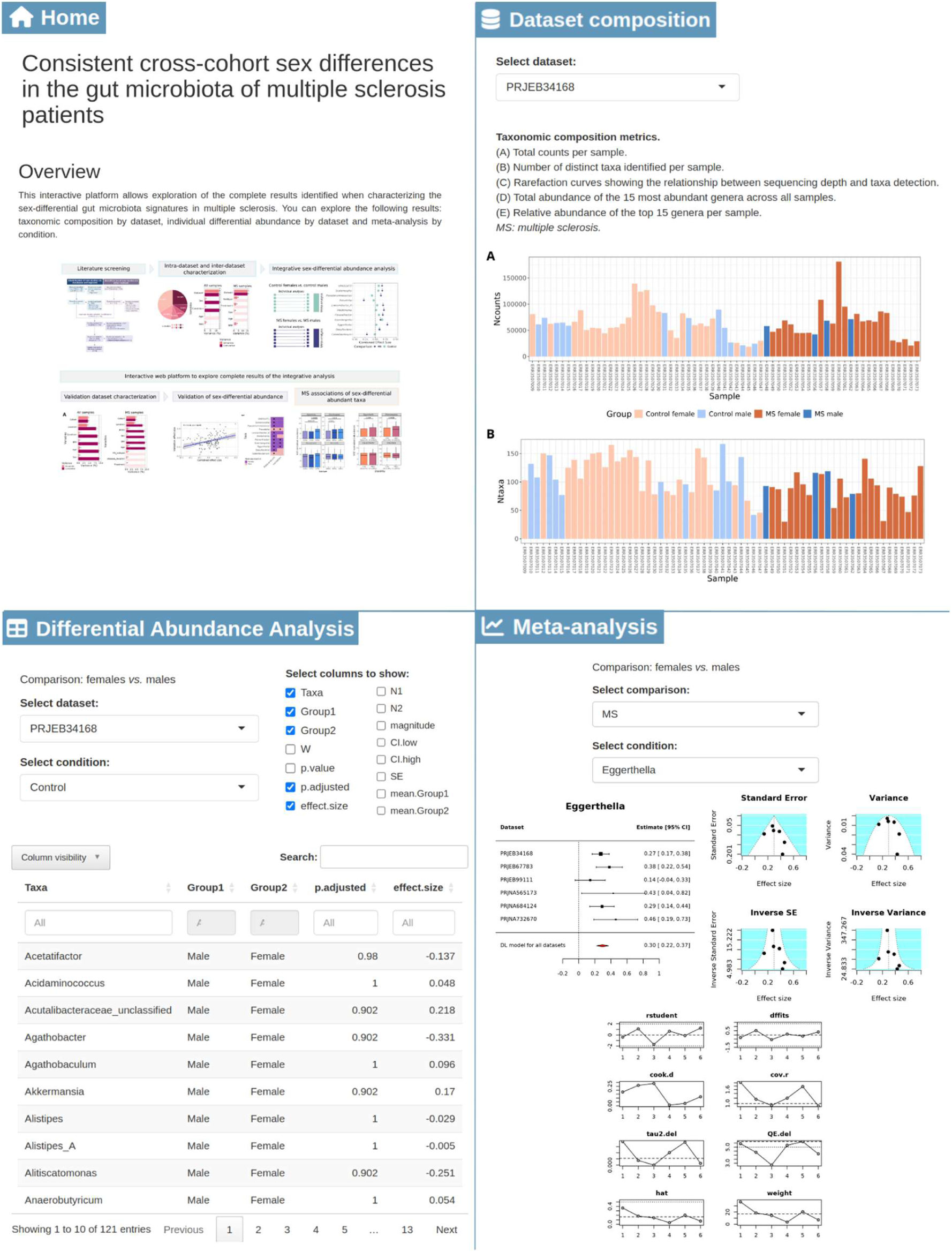
Overview of the interactive web platform. The web resource comprises four tabs: *Home*, summarizing the study design; *Dataset Composition*, reporting taxonomic composition metrics; *Differential Abundance Analysis*, presenting sex-differential results in control and MS populations for each dataset; and *Meta-analysis*, providing forest, funnel, and influence plots along with summary tables for each taxon.

## DISCUSSION

The involvement of the gut microbiota in MS pathogenesis is well established, as this disease does not develop in germ-free mouse models (4, 5), and distinct taxa have been linked to MS in human studies (24, 25). Likewise, sex is known to influence both disease prevalence and progression (26), yet sex-associated differences in gut microbiota composition in MS remains poorly characterized.

In this study, we applied an integrative approach analyzing 16S rRNA gene sequencing datasets to investigate sex-associated differences in the gut microbiota of MS patients. We observed substantial variability across studies, with the dataset of origin identified as the main source of variation (dbRDA analysis, variance >10%). Such heterogeneity has been previously reported, as technical differences including DNA extraction methods and targeted 16S rRNA gene regions can affect the microbial composition of the dataset (43–45). Beyond technical factors, this dataset-driven variability may also reflect biological heterogeneity among individuals and environmental factors (46). Specifically in MS, research findings have been discordant across studies identifying taxa that differ between MS patients and healthy controls. To overcome this, systematic reviews (22, 23) and meta-analyses (24, 25) have sought to identify consensus patterns.

Within this context, we conducted sex-differential abundance analyses to characterize dataset-specific shifts. The results were then integrated using a random-effects meta-analysis strategy. Sex-associated differences in MS (i.e., MS females *vs.* MS males) were more pronounced and consistent than in controls (i.e., control females *vs.* control males). Particularly, we identified 11 taxa with significant differences in MS, nine more abundant in females and two more abundant in males. These findings were validated using the iMSMS consortium dataset (15), the largest and most geographically diverse cohort identified in our systematic review. Overall, nine of the eleven patterns were replicated, and four remained significant after multiple testing correction: *Eggerthella*, *Eisenbergiella*, and *Flavonifractor* were more abundant in MS females, whereas *Prevotella* was more abundant in MS males. Notably, *Eggerthella* and *Eisenbergiella* were also associated with specific MS subtypes and higher disability degree.

*Eggerthella* has been associated with pro-inflammatory environments in both mouse models (47) and humans (48). It has been quantitatively linked to the human enterotype associated with systemic inflammation (Bact2), increasing in abundance in overall lower microbial load fecal microbiota (49). Moreover, Chen *et al.* (49) identified *Eggerthella* as one of the top taxa enriched in rheumatoid arthritis, where this bacteria may exacerbate disease severity. The connection of *Eggerthella* to immune-mediated diseases may involve the activation of Th17 cells (50, 51), which play a pivotal role in MS pathogenesis to trigger autoimmunity (52). Indeed, *Eggerthella* has been reported to be more abundant in MS patients compared with healthy controls (6). Our observations further suggest a higher abundance in females, that may reflect either its contribution to the enhanced inflammation observed in females versus males, or reflect its fitness advantage in the autoimmune context. We additionally found that *Eggerthella* abundance was positively associated with disease duration - but not with age in controls - and with the SPMS subtype, suggesting a potential link with MS chronicity.

*Eisenbergiella* was first reported in 2014 as a genus with the capacity for short-chain fatty acids (SCFAs) production (53). *Eisenbergiella* has been reported to be enriched in diverse conditions, like gestational diabetes mellitus (54), end-stage renal disease (55), colorectal cancer (56) and the autoimmune disease systemic lupus erythematosus (57). It was also associated with neurodegenerative diseases, reported as potentially protective in Alzheimer’s disease (58) but enriched in Parkinson’s disease (59). In MS-based studies, it was identified as more abundant when compared with healthy controls, both in human cohorts and in EAE models (15, 60). Yoon *et al.* (60) also reported that ileal microbiota derived from patients with MS, but not from their healthy monozygotic twins, induced MS-like disease in germ-free transgenic mice. Within this context, *Eisenbergiella* emerged as a candidate disease-associated taxon, suggesting a potential contributory role in MS pathogenesis, to which we now add a sex-differencial role being female-enriched. We also observed a positive correlation with age in MS and controls, consistent with previous reports linking *Eisenbergiella* abundance to long-lived individuals (61).

Although we did not identify direct associations between these taxa and MS clinical features, *Flavonifractor* and *Prevotella* are known to modulate the host immune system. *Flavonifractor* was identified as more abundant in females with MS compared with males with MS. This genus is involved in the metabolism of dietary flavonoids, bioactive compounds derived from fruits and vegetables (62). *Flavonifractor* has been associated with colorectal cancer, authors suggesting a potential role for its metabolic ability to degrade anticarcinogenic flavonoids (63). However, experimental studies in mice suggest a protective role, contributing to the attenuation of autoimmune responses (64, 65) and to improved responses to CAR-T therapy (66).

*Prevotella* was the single validated taxon more abundant in males than in females with MS. This genus has been linked to diets rich in fiber and complex carbohydrates (67). Of note, in the meta-analysis from Qingqi Lin *et al.* (24) this taxon was reported to be decreased in MS patients compared with controls across all included studies, reaching statistical significance in more than half. *Prevotella* has been shown to suppress Th17-mediated autoimmune responses and to reduce disease severity in EAE (68). Nevertheless, it has also been associated with chronic inflammation (69), suggesting that its immunomodulatory role is highly context-dependent. Our results suggest that studying the role of these taxa in the pathophysiology of MS using experimental models should be done with sex-effects kept in mind.

To the best of our knowledge, this manuscript describes the first integrative study that explicitly evaluates the influence of sex in MS within the context of the human gut microbiome. Some of the identified taxa had previously been associated with MS without considering the sex differences. Notably, some of the validated taxa show comparable sex-associated patterns in other diseases, such as *Eggerthella*, which has been reported to be more abundant in females in metabolic syndrome (70) and bronchial asthma (71). These convergent observations suggest the existence of shared, sex-related mechanisms across conditions. In addition, we identified microbial associations with clinical characteristics of MS, contributing to the characterization of how sex-specific microbiota composition may relate to disease course.

Limitations in this study should be acknowledged. Heterogeneity across cohorts, including differences in sample size, sequencing platforms, and targeted 16S rRNA regions, represents an important source of variability that may limit the generalizability of our findings. Limited metadata availability across studies restricted our ability to account for potential confounders beyond disease status, sex, BMI, MS treatment, and age. Microbial cell counts required for quantitative profiling were also unavailable, limiting our analyses to relative microbial abundances. Furthermore, sex-related differences in gut microbiota, while more pronounced in patients, appeared as non-significant trends in the controls. These results suggest that sex-differences should be explored in larger cohorts of healthy controls to further understand the extent and variation in gut microbiota sex-differences within health boundaries. Finally, as this work relied on 16S rRNA gene sequencing, future studies using whole-genome shotgun metagenomics and metabolomics will be essential to elucidate functional and mechanistic links between sex, gut microbiota, and MS.

Overall, despite the inherent challenges of microbiome research, this study provides a novel and integrative perspective on the interplay between sex, the gut microbiota and MS. Results are readily accessible through a freely available, user-friendly web resource to facilitate further research. Sex-differential microbial composition may have the potential to influence MS pathogenesis, underscoring the need for further research to clarify their role in disease modulation.

## METHODS

Bioinformatics analyses were performed using a combination of bash scripting and R programming (v.4.3.2). The developed code is available at https://github.com/IrSoler/cbl-metaanalysis-16SMS.

### Systematic review

A systematic review was conducted according to the Preferred Reporting Items for Systematic Reviews and Meta-Analyses (PRISMA) 2020 standards (72). Responses to the PRISMA abstract checklist are provided in the **Supplementary Material**. Literature screening was performed up to April 2025 using the European Nucleotide Archive (ENA) and the Sequence Read Archive (SRA) public repositories. Peer-reviewed publications with associated data were also searched for in PubMed. The following queries were applied: for ENA, *multiple sclerosis microbiome*; for SRA, *((multiple sclerosis) AND “human gut metagenome”[orgn: txid408170]) AND bioproject_sra[filter]*; and for PubMed *(“multiple sclerosis”[All Fields] AND ((micro*[All Fields]) AND (human[All Fields] OR “homo sapiens”[All Fields]) AND (metagenom*[All Fields] OR metatranscript*[All Fields] OR 16S[All Fields])))*.

All identified studies were individually reviewed. Studies were excluded if they met any of the following criteria: i) Methodology: studies that did not generate 16S metagenomic data, ii) Sample origin: studies not based on human fecal samples, iii) Disease examined: studies not focused on MS, iv) Experimental design: studies including only MS patients without control samples, v) Sex information: studies that did not record sex information, vi) Sample size: datasets with fewer than three individuals per condition and sex, vii) Data availability: unavailability of FASTQ files and/or associated metadata.

FASTQ files and associated metadata from the selected studies were retrieved for primary bioinformatic analysis. Metadata nomenclature was standardized across all studies.

### Data processing for individual studies

Raw sequencing reads were filtered and trimmed using Cutadapt (73) (v.4.6) to remove low-quality bases and/or reads, adapter sequences (when present), and undetermined nucleotides. Detailed quality control thresholds for each dataset are provided in **Supplementary Table 2**. Amplicon Sequence Variants (ASVs) were inferred using the DADA2 pipeline (74) (v.1.30.0). Briefly, the workflow included dereplication, error rate learning, ASV inference, merging of paired-end reads (when applicable), chimera removal, statistical evaluation, construction of the ASV abundance matrix, and filtering of sequences with non-target lengths. Predefined parameters were applied, except for *minFoldParentOverAbundance* used for chimera detection, which was set to 8. Reads were discarded if they were not identified as ASVs, if forward and reverse reads could not be successfully merged (in paired-end datasets), if they were identified as chimeras, or if their length fell outside the expected range for the amplified region.

Taxonomic assignment was performed using the Genome Taxonomy Database (GTDB) release 220 (75). Taxonomic annotations for direct use in DADA2 were obtained from Dr. Claus Christophersen’s team processed files (76). Each ASV was taxonomically classified at *Kingdom*, *Phylum*, *Class*, *Order*, *Family*, and *Genus* levels. ASVs that could not be confidently classified at a given taxonomic level were assigned as *unclassified* at the last reliably determined rank.

Quality assessment was performed to exclude unsuitable samples for downstream analyses. Samples were removed if they met any of the following exclusion criteria: (i) total read counts below 1,000, (ii) fewer than 30 distinct taxa identified, (iii) rarefaction curves not reaching a clear *plateau*, or (iv) evidence of contamination or poor processing based on the dominant taxonomic composition. The remaining samples constituted the filtered abundance matrices, which were the input for subsequent analyses that were performed at the genus level.

### Characterization of microbiota diversity

Inter- and intra-study variability were characterized by quantifying within-sample diversity (alpha diversity, Shannon index) and between-sample dissimilarity (beta diversity, Bray-Curtis dissimilarity metric) with functions implemented in the phyloseq R package (77) (v.1.46.0). Multivariate associations between microbiota composition and explanatory variables were assessed by distance-based redundancy analysis (dbRDA) with the vegan R package (78) (v.2.6.8). Statistical significance was evaluated with the permutation-based ANOVA approach. Stepwise model building was performed to distinguish non-redundant sources of variation. Benjamini–Hochberg (BH) method (79) was used to correct for multiple testing, considering significance when false discovery rate (FDR) < 0.05.

### Individual differential abundance analysis

For each dataset, previously filtered abundance matrices were subset to retain genera with a relative abundance ≥ 0.01% in a minimum of 10% of the samples, reducing noise from rare or spurious taxa. Only taxa common to all datasets were selected to avoid dataset-specific biases in taxonomic representation.

Filtered matrices were individually normalized using the variance stabilizing transformation (VST) method from the DESeq2 R package (80) (v.1.42.1). Each dataset was then subjected to differential abundance testing, evaluating the impact of sex in i) the control group (*control females vs. control males*) and ii) the MS group (*MS females vs. MS males*). For each comparison, the non-parametric Wilcoxon rank-sum test was used, after confirming that no unwanted sources of variation were significant in the dbRDA analysis or the principal coordinates sample distribution.

For all studies, resulting p-values were adjusted for multiple testing using the BH method (79), with significance set at FDR < 0.05. To provide a quantitative estimate of the magnitude of observed differences, effect sizes were calculated using the *wilcox_effsize* function form the rstatix R package (81) (v.0.7.2). 95% confidence intervals for the effect sizes were estimated via bootstrap resampling with 1,000 iterations using the boot R package (82) (v.1.3-28.1).

### Meta-analysis

Meta-analysis was performed for: i) *MS females vs. MS males*, and ii) *control females vs. control males*. For each comparison, individual effect sizes were integrated with the DerSimonian-Laird method (83) using the metafor R package (84). P-values were adjusted with the BH method (79) (v.4.8.0), considering significant results when FDR < 0.05.

For each taxon and comparison, the heterogeneity indicators QE, QEp, SE, τ², I², and H² were calculated. Heterogeneity was also evaluated at study level through influence and funnel plots. For influence plots, the metrics *rstudent*, *dffits*, *cook.d*, *cov.r*, *tau2.del*, *QE.del*, *hat*, and *weight* were calculated by sequentially removing one study and assessing the impact on the model parameters. For funnel plots, standard error and variance were used as measures of variability.

### Validation of meta-analysis findings

The validation dataset (ID: PRJEB32762) was processed using the same pipeline as the datasets included in the integrative analysis. Detailed descriptions of the procedures can be found in the sections *Data processing for individual studies*, *Characterization of microbiota diversity*, and *Individual differential abundance analysis*.

In this dataset, a notable batch effect was identified associated with the enrollment group and the city where samples were collected. A combined batch variable was created by merging these two factors. Differential abundance testing was then performed using a blocked Wilcoxon rank-sum test with the *wilcox_test* function from the R coin package (85) (v.1.4.3). The formula was determined as *y ∼ x | block*, where *y* represents the abundance of the genus being tested, *x* represents the grouping variable defining the comparison of interest, and *block* corresponds to the combined batch factor.

Finally, differential abundance results were compared with those obtained from the meta-analysis considering statistical significance and direction of change.

### Microbial association with MS features

Microbial associations with MS features were evaluated using the *χ² Goodness of fit* test for categorical variables, the *Wilcoxon rank-sum* test for numerical *vs.* binary categorical variables, the *Kruskal–Wallis* tests followed by *Dunn’s post hoc* test for numerical variables across multiple groups, and *Spearman*’s correlation for numerical variables. This framework was also applied to MS features to assess potential confounding. All tests were performed with functions from stats (86) (v.4.3.2) and dunn.test (87) (v.1.3.6) R packages. Significance was considered at *p* < 0.05, or FDR < 0.05 when multiple testing was applicable.

The same approach was used to examine associations among MS features and to identify potential confounding factors between sex and other metadata variables.

### Web tool

A web tool was developed to enable interactive exploration and visualization of the individual differential abundance and meta-analysis results (https://irsoler.shinyapps.io/metaanalisis_16S_MS/). This web-based application was built using the Shiny R package (88) (v.1.9.1). It is hosted on shinyapps.io, a cloud-based platform for sharing interactive web applications built with Shiny.

## DECLARATIONS

### Ethics approval and consent to participate

Not applicable.

### Consent for publication

Not applicable.

### Availability of data and materials

The datasets analyzed in this work are publicly available: PRJNA684124 (PMID:34206853), PRJNA732670 (PMID:35472144), PRJEB99111 (PMID:28893978), PRJEB34168 (PMID:32743517), PRJNA565173 (PMID:31888483), PRJEB67783 (PMID:39402079) and PRJEB32762 (PMID:36113426). The bioinformatic code can be found at https://github.com/IrSoler/cbl-metaanalysis-16SMS and the complete results can be explored at https://irsoler.shinyapps.io/metaanalisis_16S_MS.

### Author contributions statement (CRediT-compliant)

Conceptualization: ISS, SVS, FGG; Data Curation: ISS; Investigation: ISS, CGR, SVS, VT, RGR, GF, FGG; Bioinformatic Analysis: ISS, RGR; Methodology: FGG, ISS, SVS; Supervision: FGG; Funding acquisition: FGG; Writing-Original Draft Preparation: ISS, CGR, SVS, VT, RGR, GF, FGG. All authors read and approved the final manuscript.

## Acknowledgements

The authors thank the Principe Felipe Research Center (CIPF) for providing access to the cluster, co-funded by European Regional Development Funds (FEDER) to the Valencian Community 2014-2020. The authors also thank Stuart P. Atkinson for reviewing the manuscript.

## Supplementary Information

Supplementary information includes the PRISMA checklist, Tables S1-S2, and Figures S1-S10.

